# Transcriptome analysis reveals the effects of transgenic expression of the Gal4 protein on normal gene expression in silkworm tissues

**DOI:** 10.1101/2020.02.17.952523

**Authors:** Tao Chen, Yan Ma, Rongpeng Liu, Tingting Tan, Lihua Huang, Hanfu Xu

## Abstract

The Gal4/upstream activating sequence(UAS) system, a well-known genetic tool, has been widely used to analyze gene function in many organisms, including the silkworm (*Bombyx mori*), a model lepidopteran insect. Several studies have suggested that Gal4 protein activation in tissues can negatively affect transgenic individuals; however, whether and to what extent the Gal4 protein affects normal endogenous gene expression have rarely been studied. Here, we analyzed the transcriptomes of transgenic silkworms expressing the Gal4 protein at high levels in both the wing disc (WD) and epidermis (EP) and investigated gene expression changes in both tissues. Overall, 24,593 genes were identified in the WD and EP libraries, and 2,025 and 2,488 were identified as significant differentially expressed genes(DEGs) in the WD and EP between the transgenic and control groups, respectively. These DEGs were further annotated by gene function classification and pathway assessment using public databases. In addition, 506 DEGs were shared (common) between both tissues. Of these, 97 genes were commonly upregulated, and 234 were commonly downregulated; many of them were annotated to be involved in metabolic processes such as “fat digestion and absorption”, “glycine, serine and threonine metabolism” and “glutathione metabolism” and in signal transduction pathways such as the “Rap1 signaling pathway”, “MAPK signaling pathway” and “Hippo signaling pathway”. Overall, this work enhances understanding of the effects of transgenic Gal4 protein expression on normal gene expression in silkworm tissues and suggests that researchers should pay attention to unexpected effects when using the Gal4/UAS system to study gene function.

## Introduction

The Gal4/upstream activating sequence(UAS) binary expression system, derived from yeast and originally developed in *Drosophila* [1–3], is a powerful genetic tool that allows manipulation of target gene expression in a spatiotemporally precise fashion. Since its first application in *Drosophila*, the Gal4/UAS system has been widely used to analyze gene function in dozens of organisms, including mice [4], zebrafish [5], *Xenopus* [6], *Bombyx mori* [7], *Arabidopsis thaliana* [8], *Triboliumcastaneum* [9], *Aedes aegypti* [10], *Anopheles stephensi* [11] and *Caenorhabditis elegans* [12]. The Gal4/UAS system has also been employed to develop novel genetic tools, such as the enhancer/gene trap system and the Q system [13–15], and it has been combined with genome editing tools for conditional manipulation of gene expression *in vivo* [16–17].

In recent decades, remarkable progress in gene function analysis has been achieved with the Gal4/UAS system. However, the fact that high protein levels of Gal4 have certain toxicity toward cells coexpressing UAS-linked target genes and Gal4 protein cannot be ignored. Although some researchers have described and developed novel Gal4/UAS systems with smaller sizes but greater transactivation efficiency than the original system [18–20], little attention has been paid to the effects of Gal4 on the normal expression of nontarget genes. To objectively clarify the functions of target genes, it is necessary to determine whether and to what extent Gal4 protein expression affects normal nontarget gene expression.

Recently, we established a transgenic silkworm line (named A4G4) that expresses the Gal4 mainly in the wing disc (WD) and epidermis (EP) under the control of the promoter of the *B. mori Actin4* gene [21]. This line is a good material for evaluation of the effects of Gal4 protein expression on normal gene expression in transgenic tissues. In this study, we conducted a comprehensive transcriptome analysis of WD and EP tissues and identified thousands of differentially expressed genes (DEGs) in both tissues. Our findings provide sufficient evidence, for the first time, that transgenic protein expression of Gal4 in silkworm tissues does affect the normal expression of nontarget genes.

## Materials and Methods

### Silkworm and sample collection

The wild-type (WT) silkworm strain *Nistari*, the A4G4 transgenic line, and the UtdTomato transgenic line harboring a UAS-linked red fluorescent protein variant (tdTomato) were maintained in our laboratory. The hatched larvae were reared at 24-28°C with fresh mulberry leaves. WT and A4G4 larvae at day5 of the fifth instar were selected, and the WDs and EPs were dissected, washed with precooled phosphate-buffered saline and used for subsequent experiments. The WD samples collected from WT and A4G4 larvae were named W5N_S1/S2/S3 and W5A_S1/S2/S3, respectively. The EP samples collected from WT and A4G4 larvae were named EP5N_S1/S2/S3 and EP5A_S1/S2/S3, respectively.

### RNA preparation and sequencing

Total RNA was extracted from the WD and EP samples using TRIzol Reagent (Ambion, USA) and examined on a NanoDrop 1000 spectrophotometer (Thermo Scientific, USA) and an Agilent Bioanalyzer 2100 system (Agilent Technologies, USA) for RNA integrity and quality. The qualified RNA samples were purified for poly-A-containing mRNA molecules using poly-T oligo-attached magnetic beads, fragmented into small pieces using divalent cations under elevated temperature and reverse transcribed using random primers. Thesecond-strand cDNA fragments were ligated with index adapters after being purified, end-repaired, and A-tailed. Suitable fragments were used as templates for PCR amplification. After quantification with a Qubit instrument, the PCR products were sequenced on a BGISEQ-500 platform at Beijing Genomics Institute (BGI, China).

### Alignment and quantification

The raw sequencing data were preprocessed using SOAPnuke software (https://github.com/BGI-flexlab/SOAPnuke) to remove reads with adaptors, reads with more than 5% unknown bases, and reads with low sequencing quality. The clean reads were mapped to the *B. mori* genome (http://sgp.dna.affrc.go.jp/KAIKObase/, ver.3.2.2) using HISAT software [22], and the transcripts were reconstructed using StringTie [23]. Subsequently, Cuffcompare (Cufflinks tools, [24]) was utilized to compare the reconstructed transcripts. The novel coding transcripts predicted by the Coding Potential Calculator (CPC) [25] were combined with gene models from KAIKObase to obtain a new reference gene set. In addition, the clean reads were aligned against the new reference gene set using Bowtie [26]. Gene expression levels were quantified by RSEM [27] and normalized using the fragments per kilobase per million mapped reads (FPKM) method [24,27]. The DEGseq method [28] was used to detect DEGs with adjusted *P* values <0.001. Genes were considered significant DEGs if their fold changes were >=2 and their adjusted probability values were <0.001.

### Gene Ontology (GO) and Kyoto Encyclopedia of Genes and Genomes (KEGG) enrichment analysis

The functions of the protein-coding genes were assigned according to the best matches derived from alignments to proteins in the National Center for Biotechnology Information (NCBI) nonredundant (Nr) protein sequence database using DIAMOND [29]. GO annotation was performed for all identified DEGs, and WEGO software [30] was used to conduct the GO functional classification. GO terms with adjusted *P* values ≤0.05 were defined as significantly enriched GO terms for the DEGs. Pathway enrichment analysis of DEGs was performed based on the KEGG database [31] with the same criteria.

### Data availability

The raw sequence reads are available from the NCBI search database (Bioproject PRJNA601576, run accessions SAMN13870652, SAMN13870653, SAMN13870654, and SAMN13870655). All relevant data are within the paper and its Supporting Information files.

## Results and Discussion

### Detection of Gal4 expression in transgenic silkworms

Before collecting the WD and EP samples, we first generated A4G4>UtdTomato transgenic silkworms by crossing A4G4 moths with UtdTomato moths (S1 Fig), and we detected the fluorescence distribution in various tissues of A4G4>UtdTomato silkworms to further confirm the locations of Gal4 protein expression. As shown in Fig 1, strong fluorescence intensity was detected in both the WDs and EPs but not in other tissues of the A4G4>UtdTomato individuals. In other words, the expression of UAS-linked tdTomato was activated by the Gal4 protein mainly in these two tissues, which clearly demonstrated that the Gal4 protein was expressed at comparatively high levels in the WDs and EPs of A4G4 transgenic silkworms. Then, WD and EP samples were collected from day-5 fifth-instar larvae of the WT and A4G4 lines and submitted to RNA extraction, library construction and DNA sequencing.

**Fig 1.**
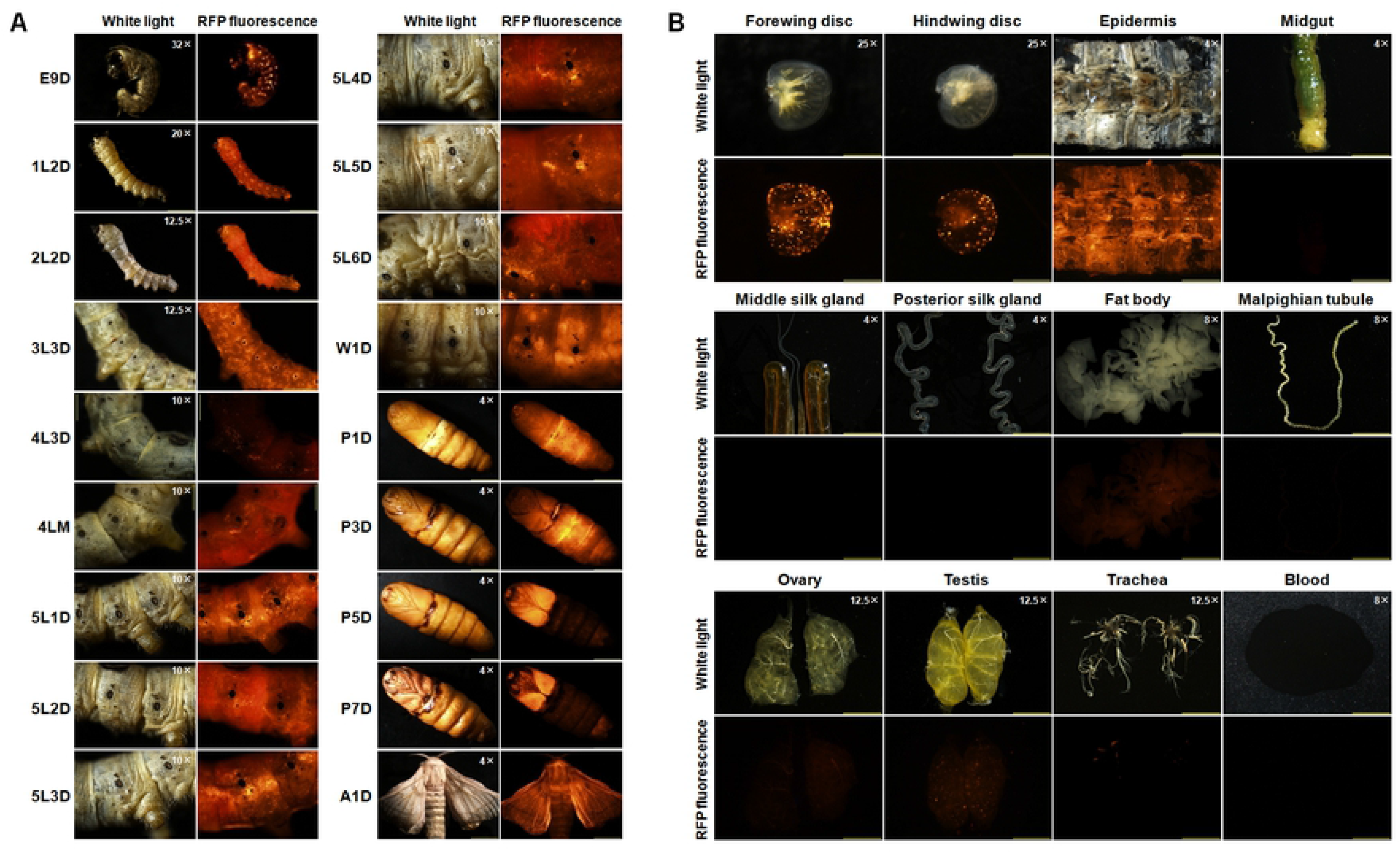
Detection of tdTomato expression in A4G4>UtdTomato transgenic silkworms. (A) Expression of tdTomato in A4G4>UtdTomato silkworms at different developmental stages. (B) Expression of tdTomato in tissues of day-5 A4G4>UtdTomato fifth-instar larvae. E9D, day-9 embryo stage; 1L2D, day-2 first-instar larval stage; 2L2D, day-2 second-instar larval stage; 3L3D, day-3 third-instar larval stage; 4L3D, day-3 fourth-instar larval stage; 4LM, fourth molting stage; 5L1D ~ 5L6D, day-1 to day-6 fifth-instar larval stages; W1D, day-1 wandering stage; P1D ~ P7D, day-1 to day-7 pupal stages; A1D, day-1 adult stage. The numbers on the photos denote the different microscope magnifications.

### Summary of the transcriptome data

RNA sequencing (RNA-Seq) generated a total of 540.19 million raw reads for all samples. After processing of these raw data, 478.36 million clean reads were obtained, with an average read depth ranging from 32.26 to 45.38 million. A total of 87.60-89.50% of the reads in each sample reached the Q30 quality score. The majority of reads in each library were mapped to the ver.3.2.2 assembly of the *B. mori* genome, and the average mapping rate of the reads was 77.57% (Table 1). The assembled transcriptome data were used to identify known and predict novel coding transcripts, which generated 24,593 genes (23,064 known genes and 1,529 novel genes). The gene number in each sample ranged from 16,439 to 18,097. Gene expression levels were calculated with RSEM software using the FPKM normalization method. Approximately 30% of the expressed genes had FPKM values larger than 10.0, and 30% had FPKM values lower than 1.0 (Fig 2).

**Table 1.** Characteristics of the RNA-Seq reads of the WD and EP samples.

| Sample | Total Raw Reads<br>(Mb) | Total Clean Reads<br>(Mb) | Clean Reads Q30<br>(%) | Clean Reads Percentage<br>(%) | Total<br>Mapped<br>(%) | Uniquely<br>Mapped<br>(%) |
| --- | --- | --- | --- | --- | --- | --- |
| W5N_S1 | 47.33 | 42.34 | 88.28 | 89.47 | 73.15 | 61.30 |
| W5N_S2 | 37.42 | 33.16 | 88.66 | 88.61 | 68.49 | 55.94 |
| W5N_S3 | 49.08 | 42.96 | 88.37 | 87.52 | 70.17 | 58.66 |
| W5A_S1 | 37.46 | 32.26 | 89.06 | 86.14 | 73.13 | 61.09 |
| W5A_S2 | 45.25 | 40.10 | 88.56 | 88.62 | 70.76 | 59.23 |
| W5A_S3 | 45.68 | 40.70 | 88.34 | 89.09 | 70.80 | 59.01 |
| EP5N_S1 | 41.18 | 36.47 | 87.70 | 88.56 | 73.26 | 60.73 |
| EP5N_S2 | 47.43 | 42.69 | 89.23 | 90.00 | 75.15 | 62.65 |
| EP5N_S3 | 49.19 | 43.92 | 89.50 | 89.28 | 76.73 | 64.15 |
| EP5A_S1 | 50.94 | 45.38 | 89.47 | 89.07 | 77.01 | 63.81 |
| EP5A_S2 | 49.19 | 43.07 | 89.22 | 87.57 | 76.59 | 63.26 |
| EP5A_S3 | 40.04 | 35.31 | 87.60 | 88.20 | 78.23 | 59.64 |

**Fig 2.**
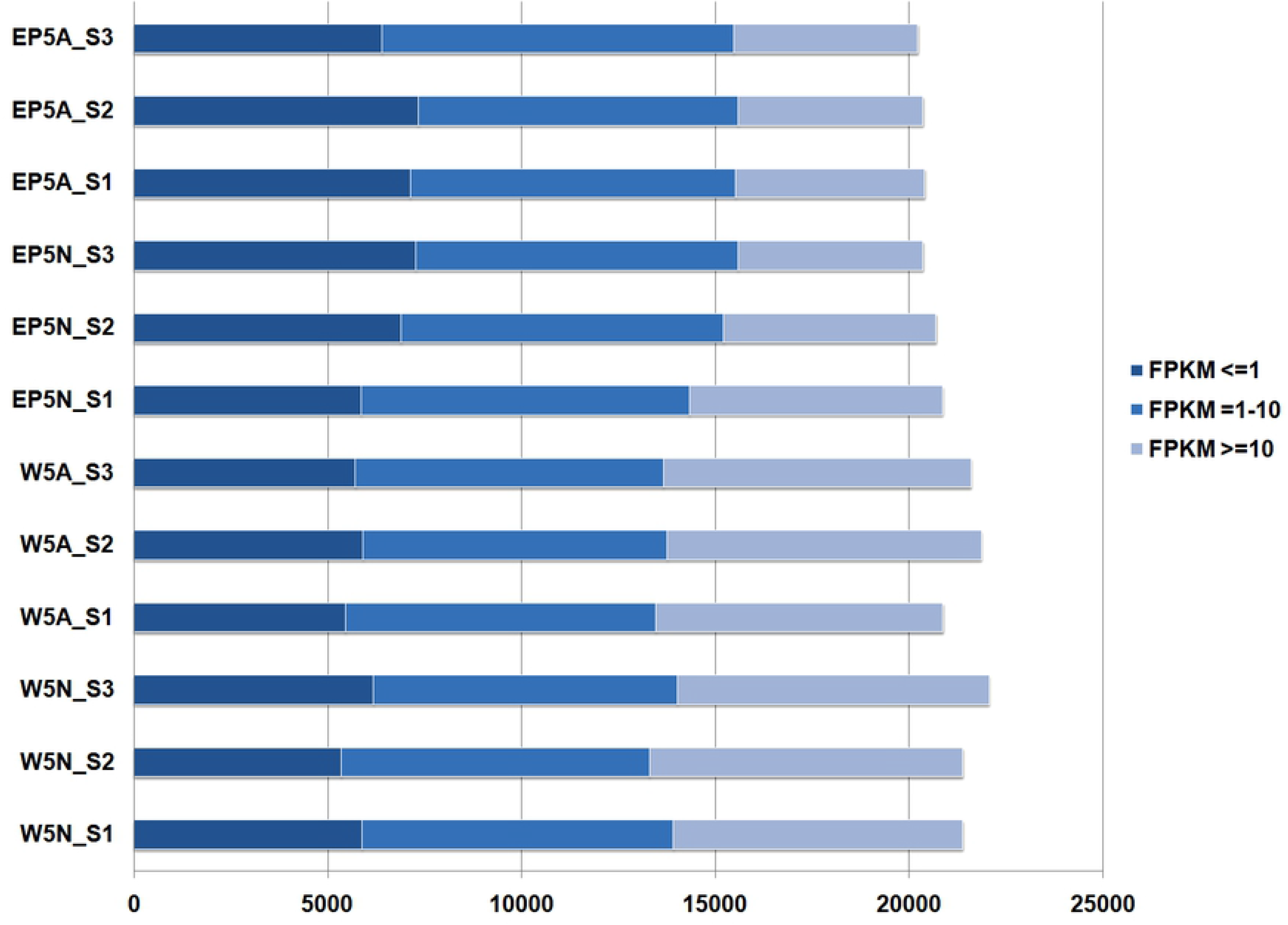
Gene expression level distribution in different FPKM value ranges. The X-axis represents the number of genes. FPKM <= 1, genes with very low expression levels; FPKM = 1-10, genes with relatively low expression levels; FPKM >= 10, genes with medium to high expression levels.

### Identification of DEGs between the transgenic and WT groups

By comparing transcriptome data between the transgenic and WT groups, a number of genes expressed in the WD and EP were identified as significant DEGs (Fig 3). In WD samples, 2,025 genes were identified as DEGs, including 771 upregulated genes and 1,254 downregulated genes (Table S1). In EP samples, a total of 2,488 DEGs were identified, among which 771 genes were upregulated and 1,717 genes were downregulated (Table S2). Thus, approximately 8.23% (2,025/24,593) and 10.12% (2,488/24,593) of the genes in the WD and EP tissues of transgenic silkworms were upregulated and downregulated, respectively, which clearly indicates that transgenic expression of the Gal4 protein in either the WD or EP affects the normal expression of endogenous genes.

**Fig 3.**
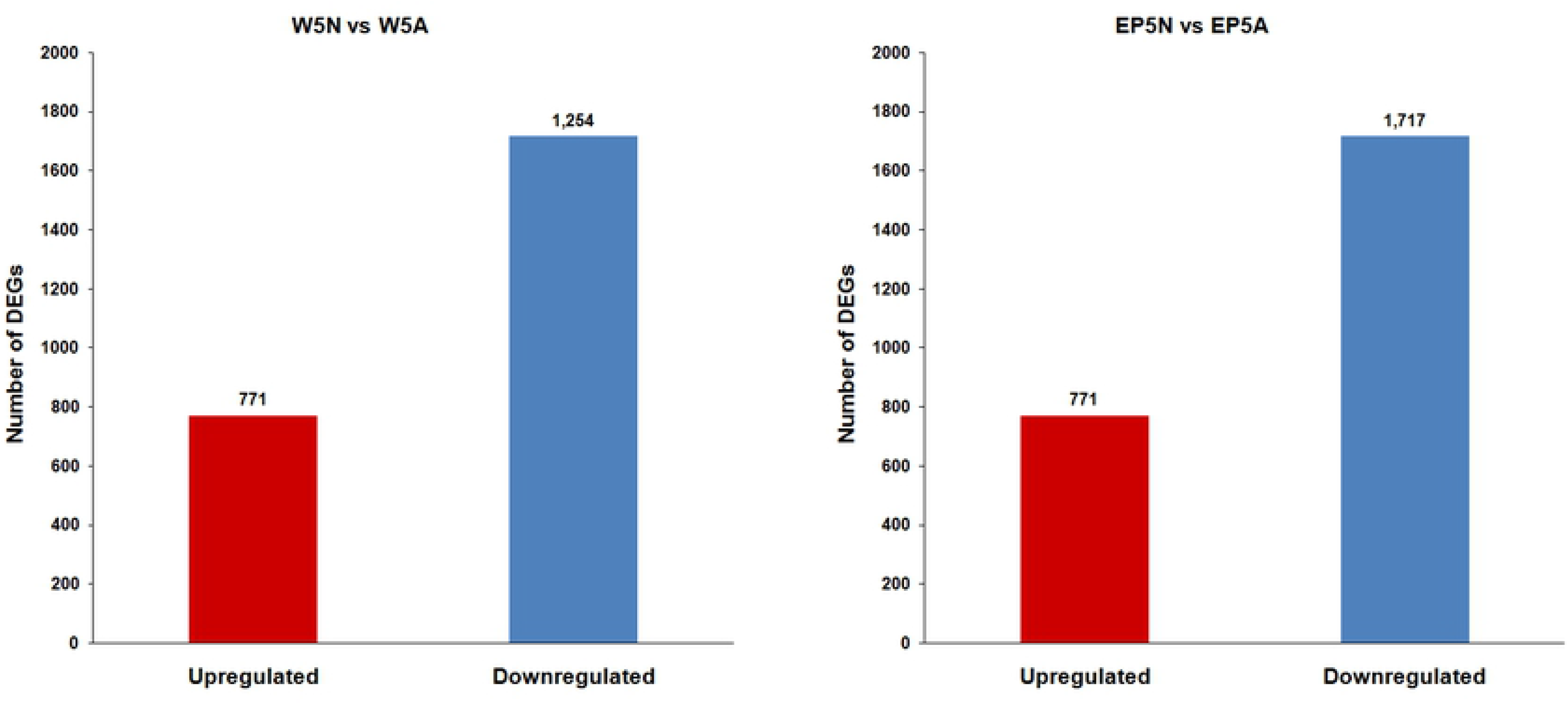
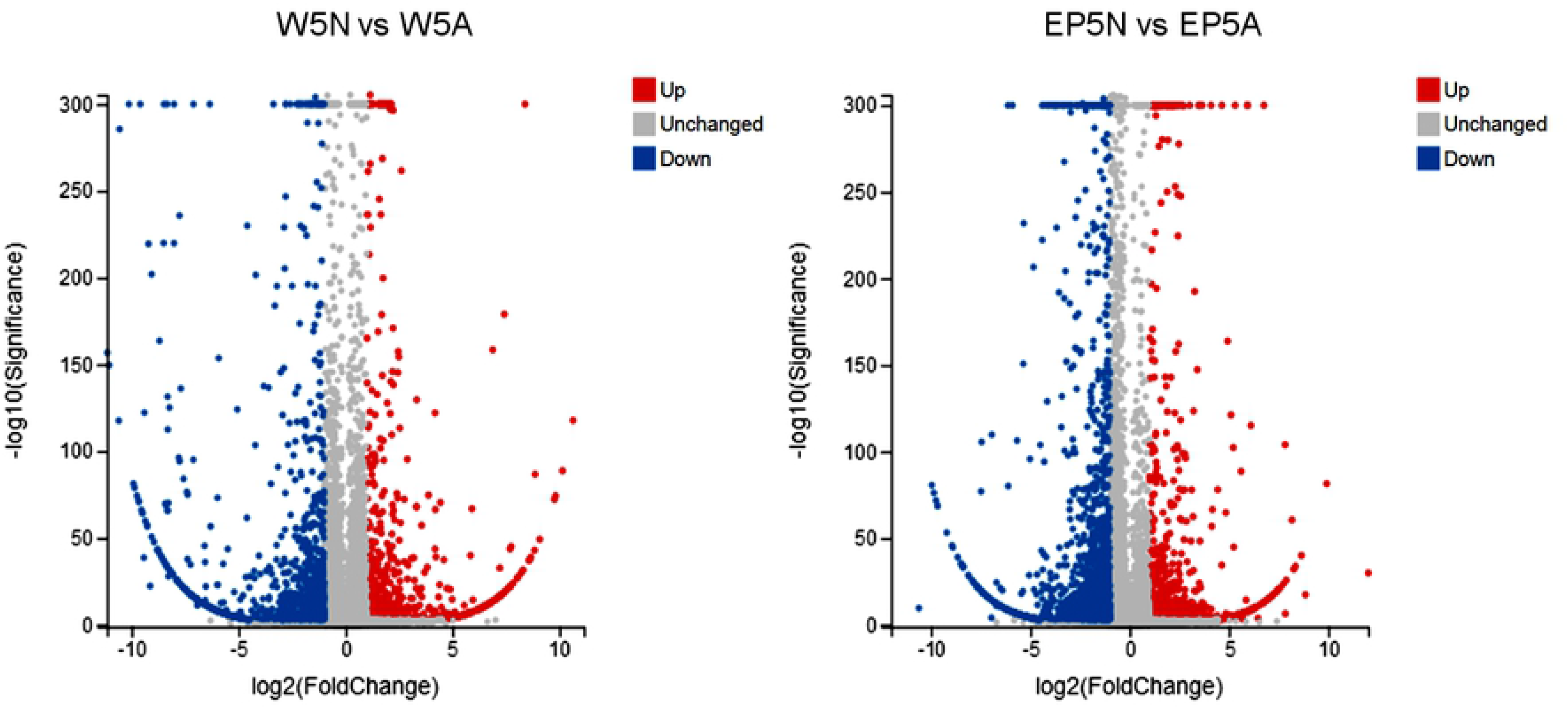
DEGs identified in the WD and EP samples. (A) Number of DEGs between the transgenic group and the WT group. (B) Volcano map of DEGs between the transgenic group and the WT group. Upregulated genes, downregulated genes, and non-significantly altered genes are indicated with red, blue, and gray points, respectively. The X-axis represents the fold change in each gene, and the Y-axis represents the significance level.

### GO annotation and KEGG pathway enrichment analysis of the DEGs

To obtain valuable information for DEG functional prediction, the DEGs were annotated with the GO database. In total, 952 DEGs in WD samples were annotated in 38 functional categories, including 15 biological process categories, 12 cellular component categories and 11 molecular function categories. Among the biological process categories, “cellular process” was the main functional group, followed by “metabolic process” and “response to stimulus”. Among the cellular component categories, “membrane” was the main functional group, followed by “membrane part” and “cell”. Among the molecular function categories, “binding” and “catalytic activity” were the two main functional groups (Fig 4A). In EP samples, 1,621 DEGs were functionally annotated with 15 biological process categories, 14 cellular component categories and 10 molecular function categories. The most enriched GO terms in the biological process category were “cellular process”, “metabolic process” and “biological regulation”. The terms “membrane”, “membrane part” and “cell” were significantly enriched at the cellular component level, and the terms “binding” and “catalytic activity” were significantly enriched at the molecular function level (Fig 4B). The top 10 up- and downregulated annotated DEGs as well as the significantly enriched GO terms for the DEGs in WD and EP samples are listed in Table S3 ~ Table S6.

**Fig 4.**
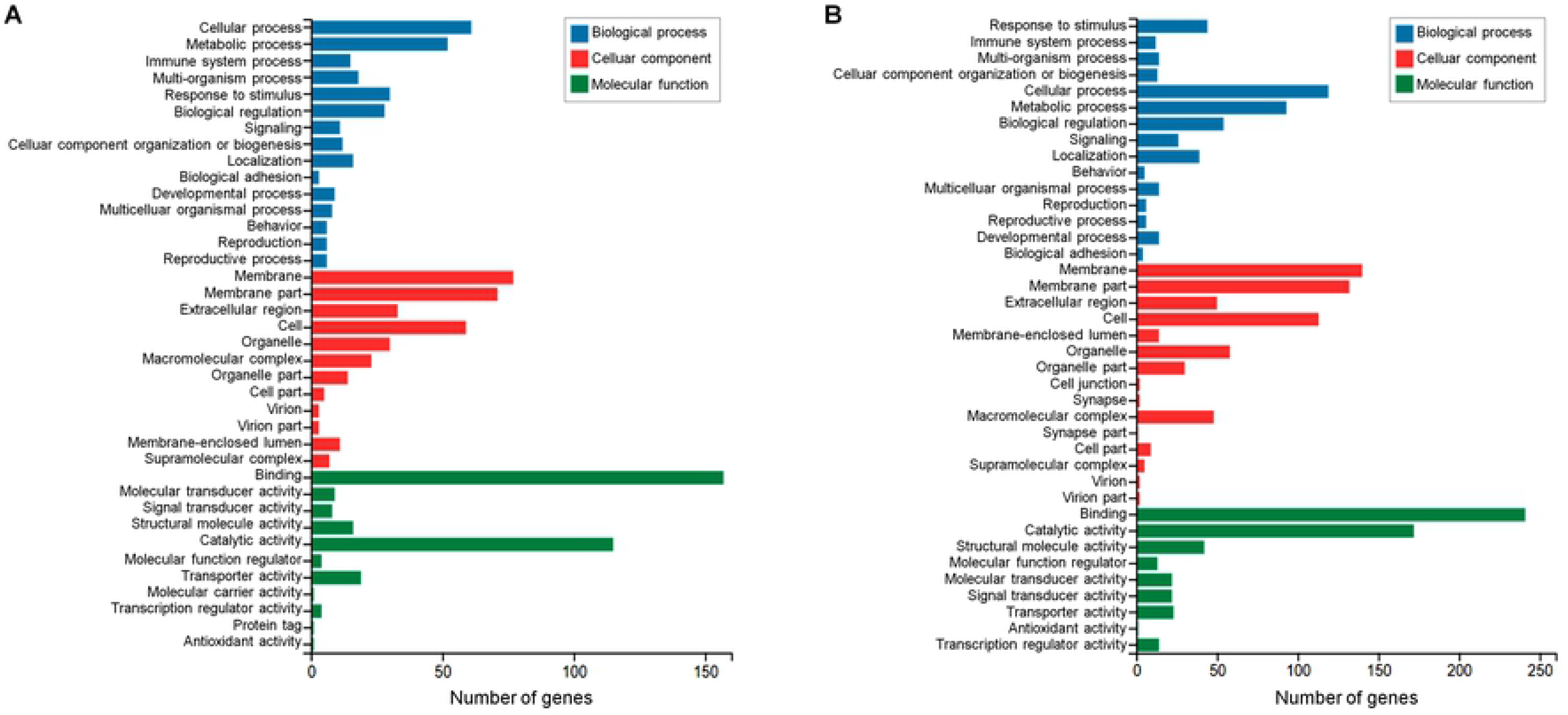
GO analysis of the DEGs between the transgenic group and the WT group. (A) GO classification of the DEGs in the WD. (B) GO classification of the DEGs in the EP.

To better interpret the pathways in which the DEGs were involved and enriched, we annotated the DEGs against the KEGG database. Briefly, the DEGs in WD samples were mainly enriched for the “phototransduction - fly”, “Influenza A”, “Hippo signaling pathway”, “fat digestion and absorption”, “viral myocarditis”, and “oxytocin signaling pathway” terms (Fig 5A). In EP samples, the DEGs were mainly annotated with the “complement and coagulation cascades”, “amoebiasis”, “tyrosine metabolism”, “ECM-receptor interaction”, “insect hormone biosynthesis”, “Hippo signaling pathway”, “axon guidance”, and “fat digestion and absorption” pathway terms, among others (Fig 5B).

**Fig 5.**
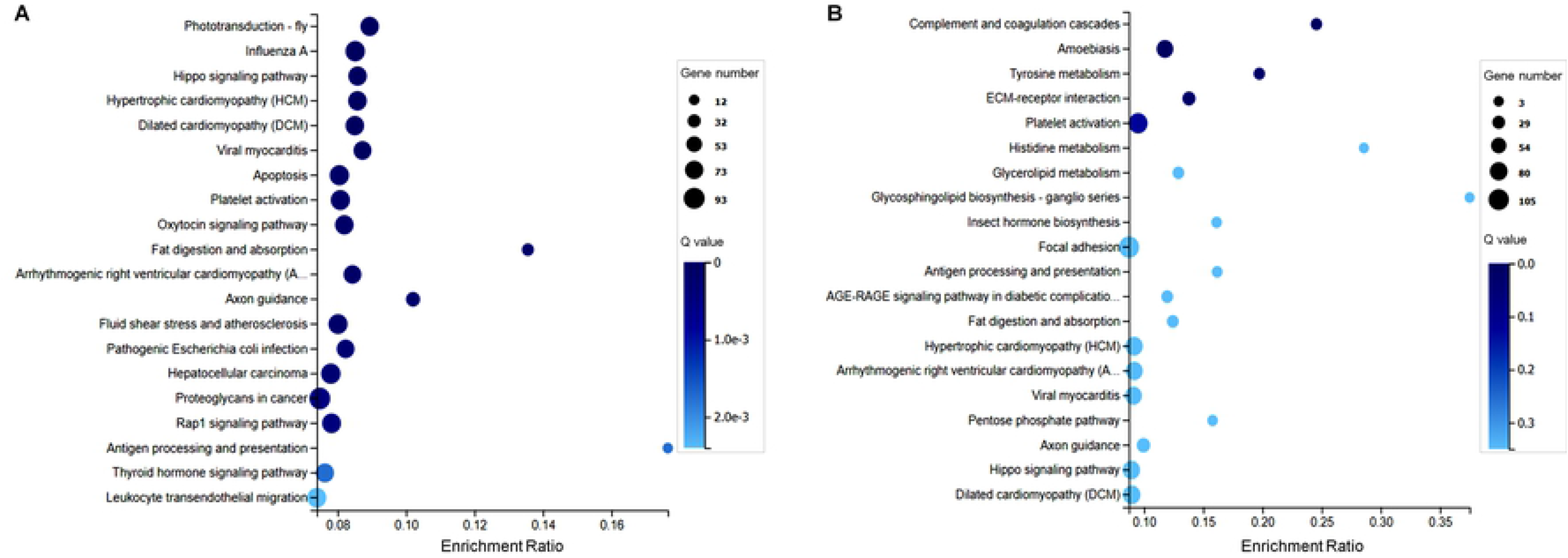
KEGG analysis of DEGs between the transgenic group and WT group. (A) KEGG classification of DEGs in the WD. (B) KEGG classification of DEGs in the EP.

### Comparative analysis of the DEGs between the WD and EP tissues

Considering that the WD and EP are known to be involved in the regulation of wing development in *B. mori*, the DEGs in both tissues were further analyzed to identify commonalities and differences. As shown in the Venn diagram in Fig 6A, 506 genes were identified as common DEGs in both tissues. Of these, 331 DEGs were common up- or downregulated genes (97 upregulated and 234 downregulated). Moreover, 111 DEGs were upregulated in WD tissues and downregulated in EP tissues, while 64 DEGs were upregulated in EP tissues and downregulated in WD tissues (Table S7). We further focused on the 331 common DEGs, the patterns of which might be influenced by the Gal4 protein in a similar way in these two tissues.

**Fig 6.**
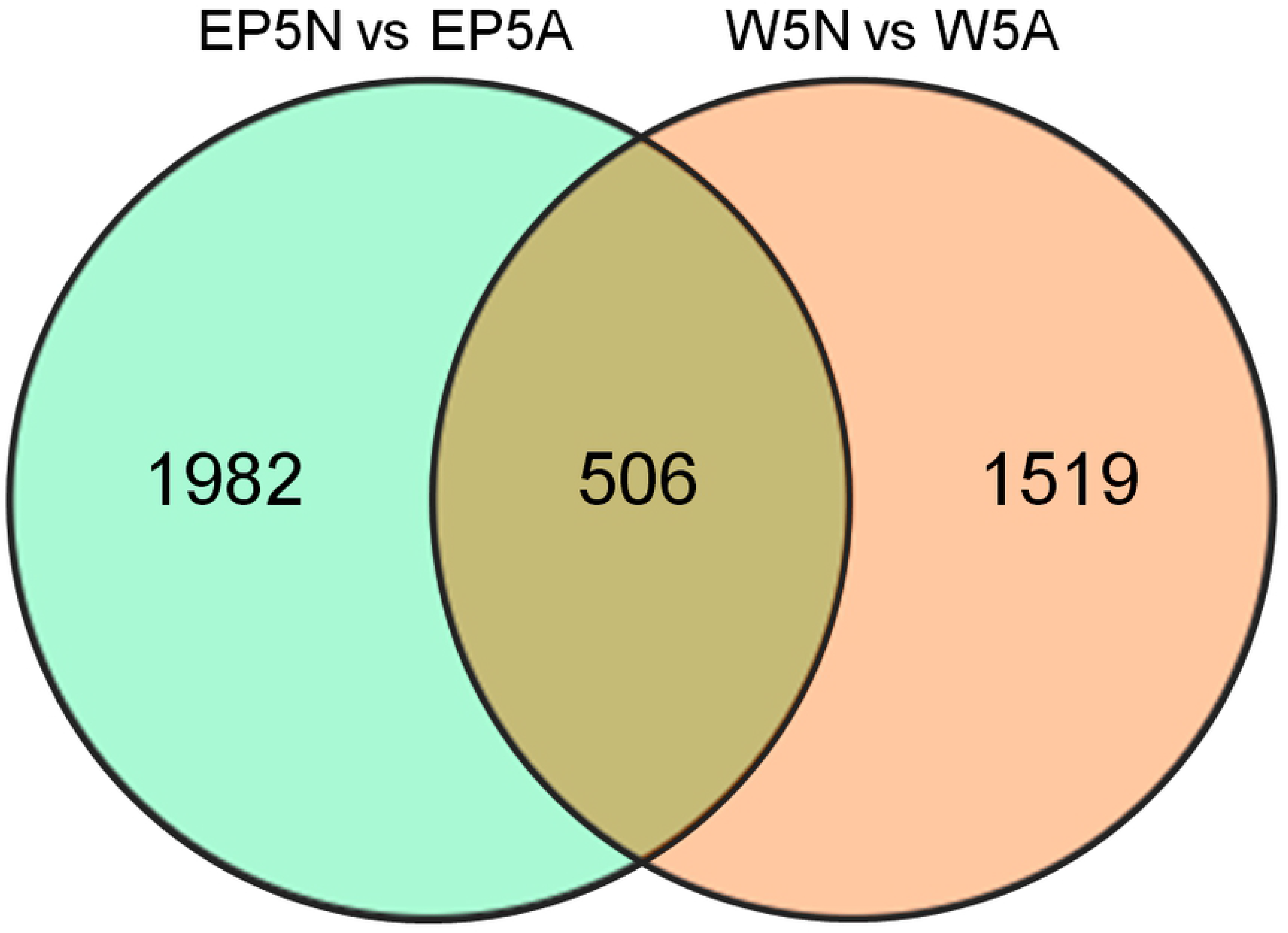
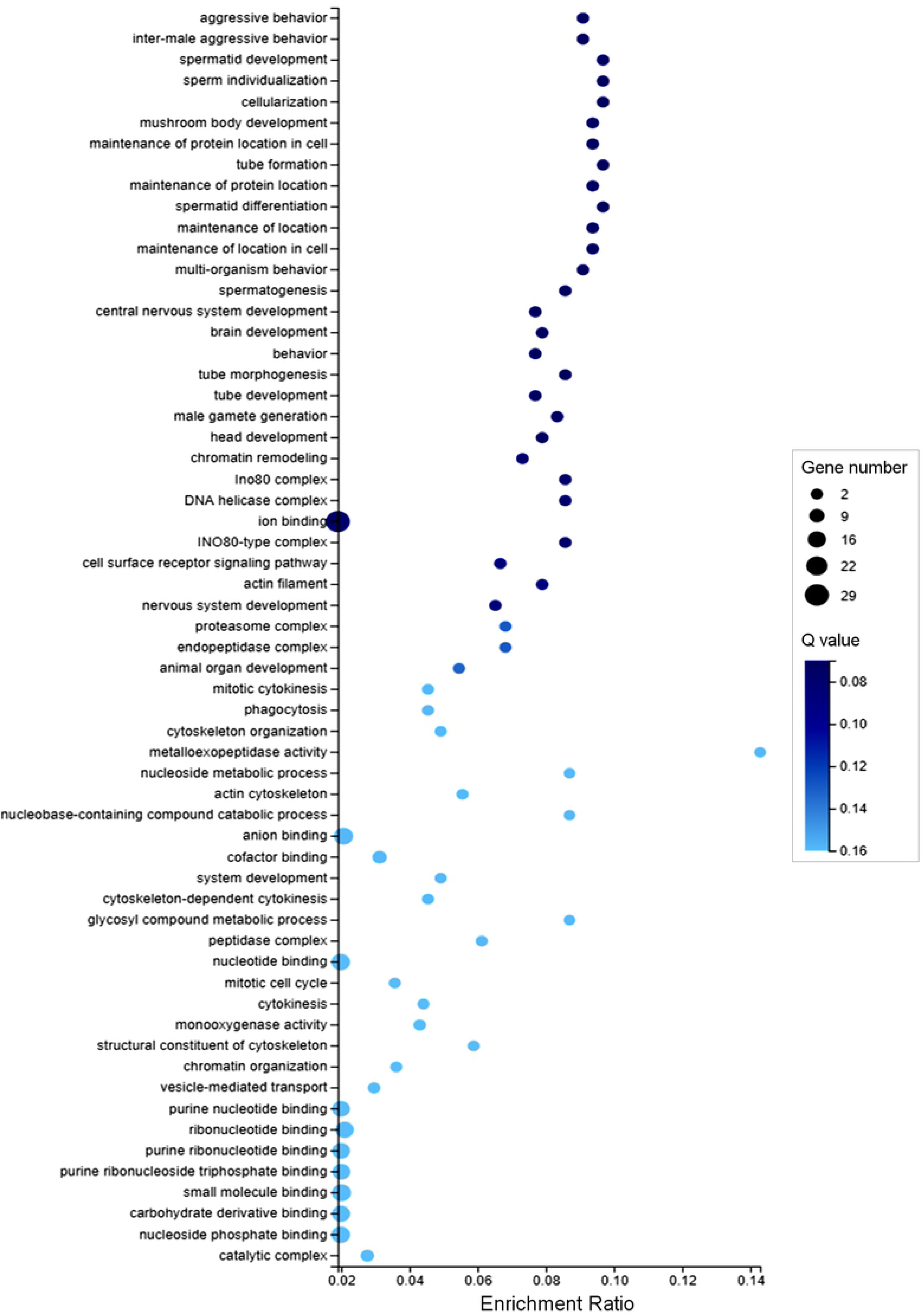
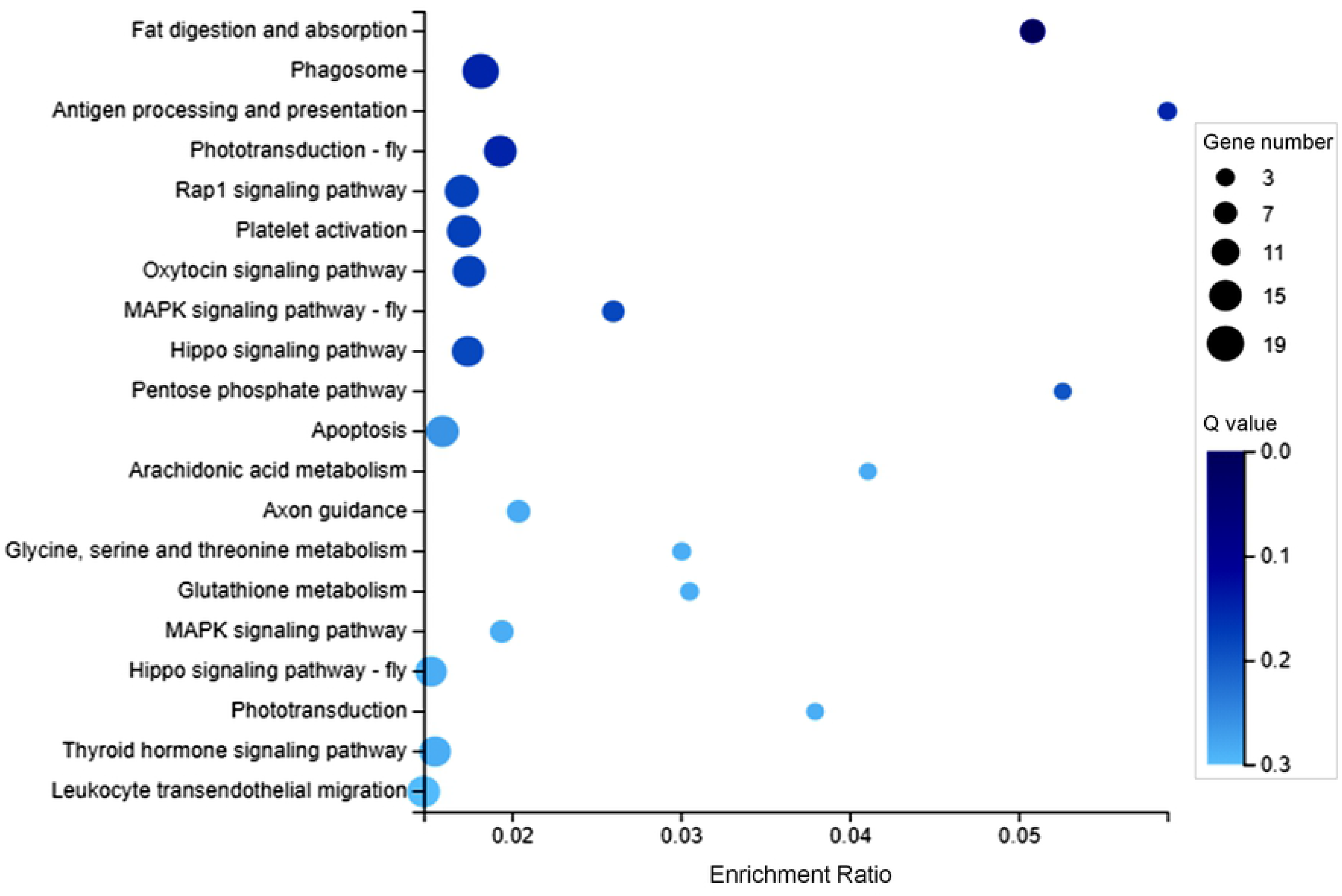
Comparison of the DEGs between WD and EP tissues. (A) Venn plot comparing DEGs in WD and EP tissues. (B) Bubble chart of the enriched GO terms for the DEGs. (C) Bubble chart of the enriched KEGG pathways for the DEGs.

First, the 331 DEGs were annotated in the NR NCBI database to a total of 208 Nr terms that encompassed 124 Nr functions. Genes annotated with more than 2 functions are listed in Table 2. The downregulated genes were annotated with the “actin-5C-like”, “actin-4”, “actin, cytoplasmic 2”, “glucose dehydrogenase”, “atlastin”, “E3 ubiquitin-protein ligase”, “26S proteasome non-ATPase regulatory subunit”, and “histidine-rich glycoprotein-like” terms. Similarly, some upregulated genes were also annotated with the “actin-5C-like”, “actin-4”, “actin, cytoplasmic 2”, “glucose dehydrogenase”, “E3 ubiquitin-protein ligase”, and “26S proteasome non-ATPase regulatory subunit” terms. These findings imply that the Gal4 protein in either the WD or EP affects the expression of actin genes as well as genes involved in metabolic processes.

**Table 2.**
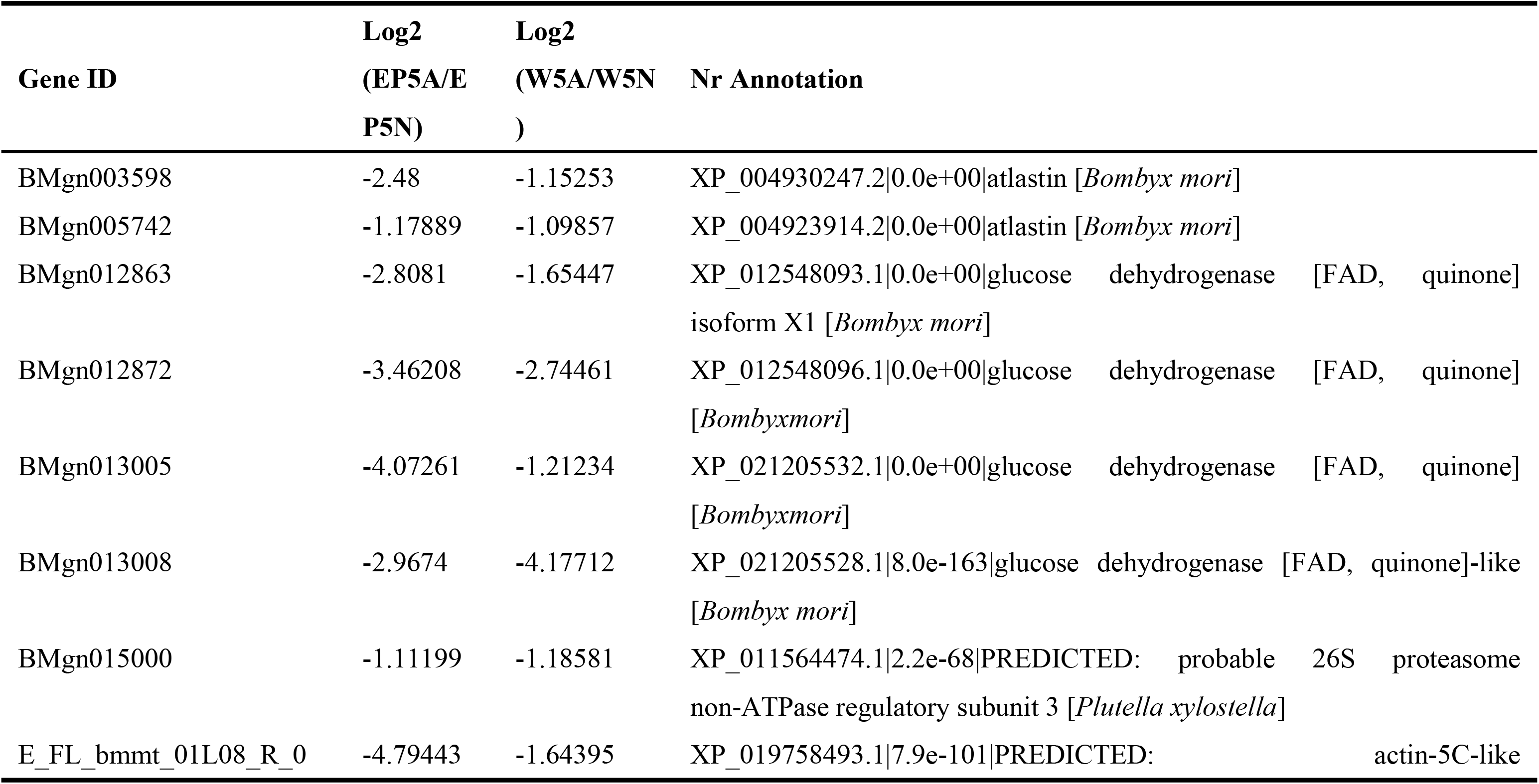
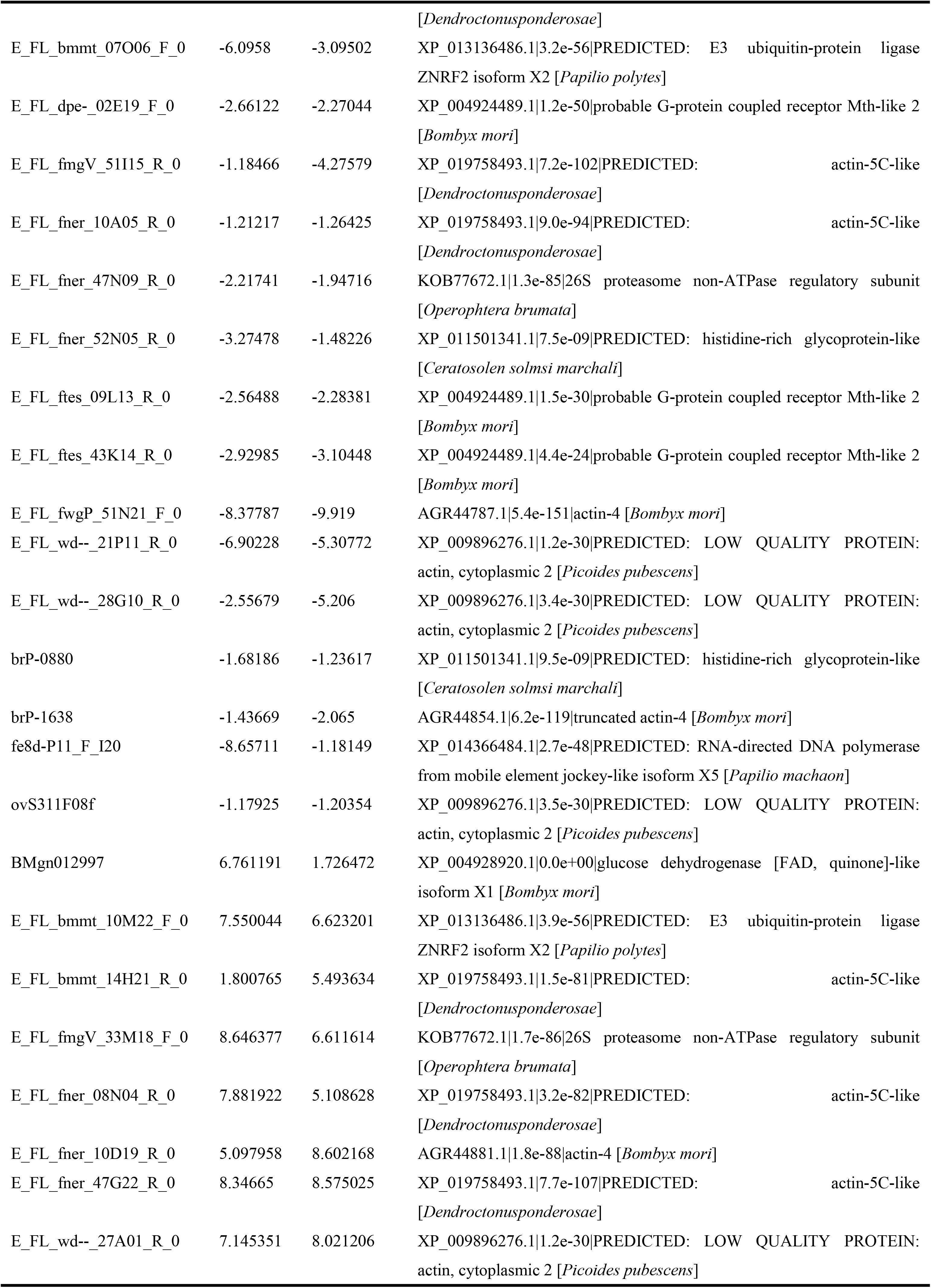
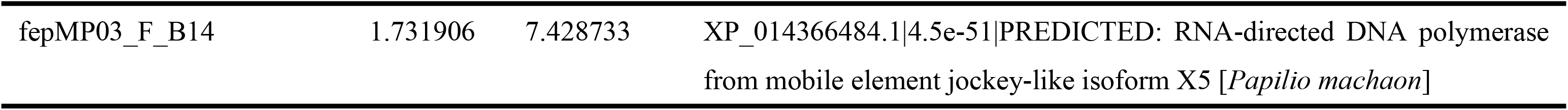
Genes annotated with more than 2 functions.

Next, we mapped all of the genes to terms in the GO database to look for significantly enriched GO terms. Among the 331 DEGs, 72 genes were annotated with 444 GO terms. Among the 60 most enriched GO terms were “multi-organism process”, “multicellular organismal process” and “biological regulation” for the biological process category; “organelle part”, “cell” and “macromolecular complex” for the cellular component category; and “binding”, “catalytic activity” and “signal transducer activity” for the molecular function category (Fig 6B). Overall, 39 common genes were annotated with these GO terms, including 16 upregulated genes and 23 downregulated genes.

Finally, the 331 DEGs were annotated against the KEGG database to better understand the biochemical pathways in which they were involved. Among the 331 DEGs,119 genes were annotated in 5 main categories. The 20 most enriched KEGG terms were “fat digestion and absorption”, “phagosome”, “antigen processing and presentation”, “phototransduction - fly”, “Rap1 signaling pathway”, “platelet activation”, “oxytocin signaling pathway”, “MAPK signaling pathway - fly”, “Hippo signaling pathway”, “pentose phosphate pathway”, “apoptosis”, “arachidonic acid metabolism”, “axon guidance”, “glycine, serine and threonine metabolism”, “glutathione metabolism”, “MAPK signaling pathway”, “Hippo signaling pathway - fly”, “phototransduction”, “thyroid hormone signaling pathway”, and “leukocyte transendothelial migration” (Fig 6C). Overall, 59 genes were annotated with these 20 KEGG terms, 36 of which were downregulated and 23 of which were upregulated. Taken together, these functional annotations suggest that the expression of genes involved in a series of metabolic processes and signal transduction pathways is influenced by the Gal4 protein in either the WDs or the EPs of transgenic silkworms.

## Conclusion

In this study, RNA-Seq, de novo assembly and functional annotation were performed to characterize the transcriptome profiles of two types of tissues, the WD and EP, that were collected from day-5 fifth-instar larvae of silkworms of the WT *Nistari* line and the transgenic A4G4 line expressing the Gal4 protein mainly in the WD and EP. We conducted comparative transcriptome analyses to identify the DEGs and potentially related biological pathways in the silkworms. A total of 2,025 and 2,488 genes were identified as significant DEGs in the WD and EP, respectively, among which 506 DEGs were common to both tissues (97 were commonly upregulated, and 234 were commonly downregulated). Many of the common genes were annotated to be involved in metabolic processes (“fat digestion and absorption”, “glycine, serine and threonine metabolism”, “glutathione metabolism”, etc.), and signal transduction pathways (“Rap1 signaling pathway”, “MAPK signaling pathway”, “Hippo signaling pathway”, etc.). Overall, our results present a comprehensive view of gene expression profiles in the WDs and EPs of WT and A4G4 silkworms and reveal that transgenic expression of the Gal4 protein affects normal gene expression in these tissues. Our findings also provide timely and valuable information for future studies on gene function using the Gal4/UAS binary system.

## Acknowledgments

This work was supported by grants (31872291) from the National Natural Science Foundation of China and grants (cstc2017jcyjBX0041) from the Chongqing Research Program of Basic Research and Frontier Technology. American Journal Experts performed English language editing in this manuscript.

## Author Contributions

Conceived and designed the experiments: HFX. Performed the experiments: RPL TTT. Analyzed the data: TC YM LHH HFX. Contributed reagents/materials/analysis tools: YM RPL. Wrote the paper: HFX LHH.

## Supporting Information

**S1 Fig. Generation of A4G4>UtdTomato transgenic silkworms**. Adults positive for UtdTomato (carrying 3×P3-EGFP) and A4G4 (carrying 3×P3-DsRed) showed only GFP and RFP fluorescence, respectively, in the compound eye. A4G4>UtdTomato adults showed both GFP and RFP fluorescence in the compound eye. 3×P3 is an artificial promoter that can drive EGFP or DsRed expression mainly in the ocelli and nervous tissues of *B. mori*.

**S1 Table. List of DEGs in the WD between the control and transgenic groups.**

**S2 Table. List of DEGs in the EP between the control and transgenic groups.**

**S3 Table. Top 10 annotated DEGs in the WD between the control and transgenic groups.**

**S4 Table. Top 10 annotated DEGs in the EP between the control and transgenic groups.**

**S5 Table. Top enriched GO terms for the DEGs in the WD between the control and transgenic groups.**

**S6 Table. Top enriched GO terms for the DEGs in the EP between the control and transgenic groups.**

**S7 Table. List of common DEGs identified in both WD and EP samples.**

## References

1. Laughon A, Gesteland RF. Primary structure of the Saccharomyces cerevisiae GAL4 gene. Molecular and Cellular Biology. 1984;4: 260–267. doi:10.1128/mcb.4.2.260

2. Giniger E, Varnum SM, Ptashne M. Specific DNA binding of GAL4, a positive regulatory protein of yeast. Cell. 1985;40: 767–774. doi:10.1016/0092-8674(85)90336-8

3. Fischer JA, Giniger E, Maniatis T, Ptashne M. GAL4 activates transcription in Drosophila. Nature. 1988;332: 853–856. doi:10.1038/332853a0

4. Ornitz DM, Moreadith RW, Leder P. Binary system for regulating transgene expression in mice: targeting int-2 gene expression with yeast GAL4/UAS control elements. Proceedings of the National Academy of Sciences. 1991;88: 698–702. doi:10.1073/pnas.88.3.698

5. Scheer N, Campos-Ortega JA. Use of the Gal4-UAS technique for targeted gene expression in the zebrafish. Mechanisms of Development. 1999;80: 153–158. doi:10.1016/s0925-4773(98)00209-3

6. Hartley KO, Nutt SL, Amaya E. Targeted gene expression in transgenic Xenopus using the binary Gal4-UAS system. Proceedings of the National Academy of Sciences. 2002;99: 1377–1382. doi:10.1073/pnas.022646899

7. Imamura M, Nakai J, Inoue S, Quan GX, Kanda T, Tamura T. Targeted gene expression using the GAL4/UAS system in the silkworm Bombyx mori. Genetics. 2003;16 : 1329–1340. doi:10.1023/B:GENE.0000003842.72339.df

8. Laplaze L, Parizot B, Baker A, Ricaud L, Martinière A, Auguy F, et al. GAL4-GFP enhancer trap lines for genetic manipulation of lateral root development in Arabidopsis thaliana. Journal of Experimental Botany. 2005;56: 2433–2442. doi:10.1093/jxb/eri236

9. Schinko JB, Weber M, Viktorinova I, Kiupakis A, Averof M, Klingler M, et al. Functionality of the GAL4/UAS system in Tribolium requires the use of endogenous core promoters. BMC Developmental Biology. 2010;10: 53. doi:10.1186/1471-213x-10-53

10. Kokoza VA, Raikhel AS. Targeted gene expression in the transgenic Aedes aegypti using the binary Gal4-UAS system. Insect Biochemistry and Molecular Biology. 2011;41: 637–644. doi:10.1016/j.ibmb.2011.04.004

11. Obrochta DA, Alford RT, Pilitt KL, Aluvihare CU, Harrell RA. piggyBac transposon remobilization and enhancer detection in Anopheles mosquitoes. Proceedings of the National Academy of Sciences. 2011;108: 16339–16344. doi:10.1073/pnas.1110628108

12. Wang H, Liu J, Gharib S, Chai CM, Schwarz EM, Pokala N, et al. cGAL, a temperature-robust GAL4–UAS system for Caenorhabditis elegans. Nature Methods. 2016;14: 145–148. doi:10.1038/nmeth.4109

13. Yang MY, Armstrong J, Vilinsky I, Strausfeld NJ, Kaiser K. Subdivision of the drosophila mushroom bodies by enhancer-trap expression patterns. Neuron. 1995;15: 45–54. doi:10.1016/0896-6273(95)90063-2

14. Asakawa K. The Tol2-mediated Gal4-UAS method for gene and enhancer trapping in zebrafish. Methods. 2009;49: 275–281. doi:10.1016/j.ymeth.2009.01.004

15. Potter CJ, Tasic B, Russler EV, Liang L, Luo L. The Q System: A Repressible Binary System for Transgene Expression, Lineage Tracing, and Mosaic Analysis. Cell. 2010;141: 536–548. doi:10.1016/j.cell.2010.02.025

16. Xue Z, Wu M, Wen K, Ren M, Long L, Zhang X, et al. CRISPR/Cas9 Mediates Efficient Conditional Mutagenesis in Drosophila. G3 (Bethesda). 2014;4: 2167–2173. doi:10.1534/g3.114.014159

17. Santis FD, Donato VD, Bene FD. Clonal analysis of gene loss of function and tissue-specific gene deletion in zebrafish via CRISPR/Cas9 technology. Methods in Cell Biology The Zebrafish-Genetics, Genomics, and Transcriptomics. 2016;135: 171–188. doi:10.1016/bs.mcb.2016.03.006

18. Köster RW, Fraser SE. Tracing Transgene Expression in Living Zebrafish Embryos. Developmental Biology. 2001;233: 329–346. doi:10.1006/dbio.2001.0242

19. Kobayashi I, Kojima K, Uchino K, Sezutsu H, Iizuka T, Tatematsu K-I, et al. An efficient binary system for gene expression in the silkworm, Bombyx mori, using GAL4 variants. Archives of Insect Biochemistry and Physiology. 2011;76: 195–210. doi:10.1002/arch.20402

20. Zhang Y, Ouyang J, Qie J, Zhang G, Liu L, Yang P. Optimization of the Gal4/UAS transgenic tools in zebrafish. Applied Microbiology and Biotechnology. 2019;103: 1789–1799. doi:10.1007/s00253-018-09591-0

21. Tan T, Liu R, Luo Q, Ma J, Ou Y, Zeng W, et al. The intronic promoter of Actin4 mediates high-level transgene expression mainly in the wing and epidermis of silkworms. Transgenic Research. 2020; doi:10.1007/s11248-020-00192-0

22. Kim D, Langmead B, Salzberg SL. HISAT: a fast spliced aligner with low memory requirements. Nature Methods. 2015;12: 357–360. doi:10.1038/nmeth.3317

23. Pertea M, Pertea GM, Antonescu CM, Chang T-C, Mendell JT, Salzberg SL. StringTie enables improved reconstruction of a transcriptome from RNA-seq reads. Nature Biotechnology. 2015;33: 290–295. doi:10.1038/nbt.3122

24. Trapnell C, Roberts A, Goff L, Pertea G, Kim D, Kelley DR, et al. Differential gene and transcript expression analysis of RNA-seq experiments with TopHat and Cufflinks. Nature Protocols. 2012;7: 562–578. doi:10.1038/nprot.2012.016

25. Kong L, Zhang Y, Ye Z-Q, Liu X-Q, Zhao S-Q, Wei L, et al. CPC: assess the protein-coding potential of transcripts using sequence features and support vector machine. Nucleic Acids Research. 2007;35. doi:10.1093/nar/gkm391

26. Langmead B, Salzberg SL. Fast gapped-read alignment with Bowtie 2. Nature Methods. 2012;9: 357–359. doi:10.1038/nmeth.1923

27. Li B, Dewey CN. RSEM: accurate transcript quantification from RNA-Seq data with or without a reference genome. BMC Bioinformatics. 2011;12. doi:10.1186/1471-2105-12-323

28. Wang L, Feng Z, Wang X, Wang X, Zhang X. DEGseq: an R package for identifying differentially expressed genes from RNA-seq data. Bioinformatics. 2009;26: 136–138. doi:10.1093/bioinformatics/btp612

29. Buchfink B, Xie C, Huson DH. Fast and sensitive protein alignment using DIAMOND. Nature Methods. 2014;12: 59–60. doi:10.1038/nmeth.3176

30. Ye J, Fang L, Zheng H, Zhang Y, Chen J, Zhang Z, et al. WEGO: a web tool for plotting GO annotations. Nucleic Acids Research. 2006;34. doi:10.1093/nar/gkl031

31. Kanehisa M, Araki M, Goto S, Hattori M, Hirakawa M, Itoh M, et al. KEGG for linking genomes to life and the environment. Nucleic Acids Research. 2007;36. doi:10.1093/nar/gkm882

